# FARMS: Framework for Animal and Robot Modeling and Simulation

**DOI:** 10.1101/2023.09.25.559130

**Authors:** Jonathan Arreguit, Shravan Tata Ramalingasetty, Simon M. Danner, Auke Ijspeert

**Author notes:** Both authors contributed equally to this research (Jonathan Arreguit, Shravan Tata Ramalingasetty). Authors’ Contact Information: Jonathan Arreguit, Simon M. Danner, Auke Ijspeert.

## Abstract

The apparent ease with which animals move hides the complexity of the neuromechanical systems underlying locomotion. This behavior remains difficult to understand and replicate, motivating sustained interest across biology, neuroscience, robotics, and computer animation. Despite extensive progress, existing approaches often focus on isolated aspects of locomotion, limiting the integration of neural control, biomechanics, and environmental interactions within a unified modeling framework. Bridging biological insight, robotic implementation, and physically based character animation therefore requires shared, extensible computational tools that promote reuse, while supporting physically grounded modeling and control across animals and robots.

Our key technical contribution is the development of FARMS (Framework for Animal and Robot Modeling and Simulation), an open-source, interdisciplinary Python framework that integrates and extends existing open-source software for 3D modeling, experiment design, optimization, and data analysis within a modular and extensible workflow. FARMS lowers the barrier for constructing and simulating whole-body animal and robot models, enabling the definition of skeletal structures, musculature, and neural circuits at multiple levels of abstraction within a unified workflow.

FARMS has already been used to model and study locomotor behaviors in a variety of animals and robots, and we demonstrate the framework’s versatility and effectiveness through a series of simulation case studies. We show that FARMS enables the reuse of model components across morphologies of different complexities, while capturing mechanical interactions between the body and diverse environments, including terrestrial and aquatic settings. This allows the study and synthesis of walking, swimming, and locomotor transitions within a single, unified framework.

## 1 Introduction

Animal locomotion results from the complex interactions between the body, the nervous system, the sensory modalities, and the environment. Each of these subsystems has functional properties that emerge only when they interact with each other [Dickinson 2000]. Focusing on individual components provides only partial insights into the system’s overall behavior, making the study of animal movement a challenging and interdisciplinary scientific endeavor.

Collecting relevant data from animal experiments is essential for testing scientific hypotheses. However, this process presents several technical and ethical challenges, which vary depending on the animal species considered. For instance, measuring ground reaction forces in small animals like *Drosophila melanogaster* is currently infeasible with available technologies [Lobato-Rios et al. 2022]. Ethical constraints further complicate data collection and, in some cases, make it impossible to conduct certain experiments. These limitations highlight the importance of careful planning and specification of required measurements to ensure the feasibility of a study and to minimize unnecessary harm to animals.

To handle these challenges, the aptitude to accurately predict the outcome of experiments and identify potential causes of failure can be crucial. Predictive models based on existing knowledge can help systematically explore how different subsystems interact, thereby advancing our understanding of animal movement without relying solely on live experiments. Researchers have developed various modeling approaches that complement experimental studies, categorized into three types: (i) theoretical or analytical models, (ii) physical models, i.e. robots that mimic the behavior of animals, and (iii) numerical models. While all approaches contribute to studying locomotion and motor control, each has its strengths and limitations. Theoretical models, such as inverted pendulum models, have provided valuable insights into the general principles of locomotion [Holmes et al. 2006]. However, they are limited in their ability to capture the complex interactions of multiple subsystems. Meanwhile, physical models like robots offer promising avenues for understanding animal movement [Ijspeert 2014], but current technologies still face challenges in replicating the full complexity of animal biomechanics, such as muscle and skeletal dynamics.

In light of these challenges, physics-based numerical simulations offer an appealing alternative. Among them, *neuromechanical simulations* are an *in silico* approach to replicating animal movement by integrating multiple components, including a physical body, neural systems, and muscle models [Edwards 2010], allowing researchers to study locomotion under controlled, virtual conditions. By replicating or predicting the behaviors observed in the real world, neuromechanical simulations can provide valuable insights into the mechanics of movement, motor control, and sensorimotor integration.

However, such simulations can require significant computational resources and extensive modeling efforts. Fortunately, advancements in hardware and software tools over recent years have made these approaches more accessible. Neuromechanical simulations cannot fully replace biological experiments or robot experiments, but they can complement both by gathering additional information [Ijspeert and Daley 2023], and help make predictions.

In this work, we present the Framework for Animals and Robots Modeling and Simulation (FARMS), specifically tailored to the needs of neuromechanical simulations and animal locomotion research. FARMS emphasizes modularity, open access and extensibility, aiming to unify current efforts and accelerate progress in understanding animal motor control. Through a series of demonstrations, we present how FARMS helps design simulations with musculoskeletal models and biological neural network models for animals and robots across different environments.

## 2 Related Work

### 2.1 Neuromechanical simulations

The idea of using neuromechanical simulations to study animal movement is well established, and we will here review relevant works with a particular focus on vertebrate simulations. In the 1990s, Karl Sims showed that diverse animal morphologies and locomotory behaviors could be systematically evolved using genetic algorithms [Sims 1994a,b]. Ekeberg [1993] and Grillner et al. [1995] integrated neural networks model of spinal cord with a mechanical model of the lamprey to study swimming and body orientation. Subsequently, Ijspeert [2001] explored how a salamander could achieve both walking and swimming patterns using a neuromechanical simulation in 2.5D physics. Yakovenko et al. [2004] used a hindlimb neuromechanical model of a cat to evaluate the contribution of sensory feedback during locomotion. Ekeberg and Pearson [2005] investigated how reflex rules could trigger a hindlimb walking patterns in the cat. Harischandra et al. [2010] extended the investigation of salamander locomotion with a 3D simulator, achieving walking with all four limbs. Geyer and Herr [2010] studied in 2010 how sensory information could be used to generate bipedal walking patterns in a simple 2D simulation. Daun-Gruhn [2011] investigated how sensory information could generate the motion of the limbs in a stick insect. Aoi and Funato [2016] explored the role of muscle synergies in sensory-motor control in humans and rats for adaptive locomotion using 2D neuromuscular simulation models. Hamlet et al. [2018] investigated how curvature-based control rule combined with sensory information could achieve optimal swimming of the lamprey. Finally, Nyakatura et al. [2019] investigated the locomotion of the extinct stem amniote Orobates pabsti by leveraging simulations as a central tool for reconstructing its plausible gaits. Their simulation-driven approach provided key insights into how Orobates may have moved, demonstrating that it likely exhibited more advanced terrestrial locomotion than previously assumed for early tetrapods.

### 2.2 Neuromechanical frameworks

Previous studies on neuromechanical simulations were typically developed separately by small research groups, and would have benefited from a shared infrastructure that could reduce the need for duplication and re-implementation of models and code from scratch. The absence of such an infrastructure not only wastes resources but also limits the potential for collaboration and reproducibility. To overcome these limitations and accelerate our understanding of animal motor control, we need a comprehensive, open-source framework that integrates the components required for neuromechanical simulations across diverse applications and research domains. More precisely, while numerous projects provide specialized interfaces for physics simulation [Coumans and Bai 2016; Lee et al. 2018; Sherman et al. 2011; Smith 2006; Todorov et al. 2012], muscle modeling [Seth et al. 2018, 2011], or neural network simulations [Goodman and Brette 2009; Hines and Carnevale 1997; Stimberg et al. 2019], a critical gap remains. Existing options remain largely confined to their respective domains, and lack a higher-level interface for seamless integration between these components. As a result, modeling and simulating neuromechanical systems continues to be cumbersome, highlighting the need for a framework that bridges these specialized tools to enable more comprehensive and accessible neuromechanical simulations.

In recent years, numerous simulation frameworks have been developed, offering valuable tools for studying motor control and biomechanics. Among these, a curated selection is presented in Table 1, which highlights key features relevant to the study of animal control. While not all of these frameworks are specifically designed for neuromechanical simulations or animal locomotion, they provide flexible capabilities that can often be adapted for such purposes. More precisely, they tend to be focused on simulating robotic systems, and tools for species-specific modeling, advanced muscle and neural models, modeling specific environments such as with fluids, and optimization or learning mechanisms for tuning controllers are missing in some frameworks or are difficult to incorporate.

**Table 1.** Overview and comparison of features from different frameworks related to neuromechanical simulations. Features from other frameworks that have been enhanced or addressed in FARMS are highlighted in red.

|  | Webots | Gazebo | OpenSim | AnimatLab | Scone | Isaac | MyoSuite | FARMS |
| --- | --- | --- | --- | --- | --- | --- | --- | --- |
| **Release** | 1996 | 2004 | 2007 | 2010 | 2019 | 2021 | 2022 | 2026 |
| **Physics engine** | ODE | Dart<br>Bullet | Simbody | Bullet<br>Vortex | Simbody<br>HyFyDy | PhysX | MuJoCo | MuJoCo<br>Bullet |
| **Programing languages** | C/C++<br>Python | C++ | C++<br>Python | C++ | Lua<br>Python | C++<br>Python | Python | Python<br>Cython |
| **Muscles** | No | No | Yes | Yes | Yes | No | Yes | Yes |
| **Neural models** | No | No | No | Yes | No | No | No | Yes |
| **Data logging** | No | Yes (ROS) | No | Yes | Yes | Yes | Yes | Yes |
| **Fluids** | Drag | Partial | No | No | No | No | No | Drag/SPH |
| **GUI/Headless** | GUI | Both | Both | GUI | GUI | Both | Both | Both |
| **Playback** | No | Rosbag | No | Yes | Yes | Yes | Yes | Blender |
| **Optimization** | No | No | No | No | Yes | Gym | Gym | Yes |
| **Modeling** | Yes | Yes | Yes | Yes | No | Yes | No | Yes |
| **Active** | Partial | Yes | Yes | No | Yes | Yes | Yes | Yes |
| **License** | Apache2.0 | Apache2.0 | Apache2.0 | BSD-3-Clause | Apache2.0 | Propri. | Apache2.0 | Apache2.0 |

Webots [Michel 2004] was adopted in previous works [Crespi and Ijspeert 2006; Nyakatura et al. 2019; Thandiackal et al. 2021] due to its ease of use. With its consolidated interface, it remains an attractive choice for introductory learning and the construction of robot models. However, its reliance on the ODE physics engine is prone to instability for complex models with many joints, as a consequence of its use of maximal coordinates for solving dynamics. Gazebo [Koenig and Howard 2004] allows selecting between different physics engines, which can better support such models, but is not ideal due to its lack of direct support for a script-oriented programming language, which limits its accessibility for users with reduced programming experience. Mainly targeting robot simulations, both Webots and Gazebo do not offer direct support for muscle models. On the other hand, OpenSim [Seth et al. 2011] is an attractive candidate for its specialized tools for biomechanical modeling, including muscle models. It is a well-known tool for developing human musculoskeletal models and facilitates reverse-engineering of movement generation through motion capture. However, OpenSim is more limited for forward dynamics simulations in terms of performance and ease of use, and there remains a lack of usage related to simulating robots.

AnimatLab [Cofer et al. 2010] is to the best of our knowledge the only prior framework to offer both muscle and neural models for a multitude of animal models. However, its lack of active development since 2016 and limitations in extensibility highlight the need for more modern alternatives. It also lacks support for fluid dynamics, essential for swimming simulations, and is designed primarily as an interactive tool with the GUI, limiting its utility for batch simulations and parameter optimization, especially in headless mode to run on computer clusters.

Built upon OpenSim, Scone [Geijtenbeek 2019] offers more efficient and stable biomechanical simulations, which is accomplished by using the closed-source HyFyDy physics engine [Geijtenbeek 2021]. Aiming towards also solving OpenSim’s performance limitations for forward dynamics simulation, MyoSuite [Caggiano et al. 2022] replaces Simbody with the now open-source MuJoCo engine and provides interfaces to OpenAI’s gym API [Brockman et al. 2016]. However, both Scone’s and MyoSuite’s applications beyond biomechanical simulations remain secondary to their primary focus areas of simulating advanced biomechanics, and extending them to support robots and fluid simulations is not within their scope.

Isaac Sim [Liang et al. 2018] stands out for its reinforcement learning capabilities with robots, as it supports running multiple robot simulations in parallel thanks to its computation on the Graphics Processing Unit (GPU). However, it currently lacks integration of muscle and neural models as well as fluid environments. While efficient GPU computation support is largely beneficial, its proprietary license makes it limiting for open-source contributions.

For a comprehensive review of simulation frameworks from a robotics perspective, refer to Collins et al. [2021].

## 3 FARMS

In contrast to the frameworks listed in Table 1, FARMS focuses on including the necessary components to simulate locomotion with musculoskeletal models, biological neural network models, and the appropriate sensors for animals and robots across different environments, thereby promoting the development of neuromechanical models. It is primarily written in Python and features a range of tools to help design models with realistic dynamics, to run controllers based on neural models, and for comparative studies with animals and robots. It also supports optimization methods for parameter tuning and 3D visualization for debugging and rendering high-quality videos for publication.

At an implementation level, FARMS leverages and integrates available open-source software into a set of implemented packages developed to facilitate interoperability between the different components and manage dependencies. For this purpose, it has been split into a series of packages to enable the users to pick specific components required for their experiments, and to facilitate development and handle contributions. Considerable effort was also put into integrating wrapper functions such that users could create and incorporate their own models and controllers and run physics-based simulations, with support for multiple physics engines, similarly to Gazebo [Koenig and Howard 2004]. For example, the farms_muscle package provides Hill-type muscle models, enabling the simulation of biomechanical systems within a simulator. Another example is the farms_network package, used by Lobato-Rios et al. [2022] for simulating sparse neural networks, which can function independently or be seamlessly integrated with FARMS. In essence, FARMS can be viewed as a basis upon which wrapper code is written to glue existing software together.

Within FARMS, developing a neuromechanical simulation can be broken down into four main components (Figure 2):

**Fig. 1.**
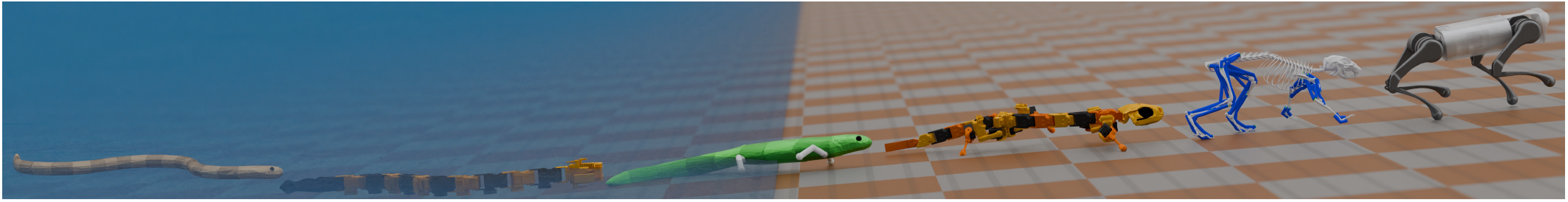
Gait snapshots of animal and robot locomotion in water, land and transition taken at specific time points from simulations within FARMS framework. Model sizes have been scaled for illustration purpose.

**Fig. 2.**
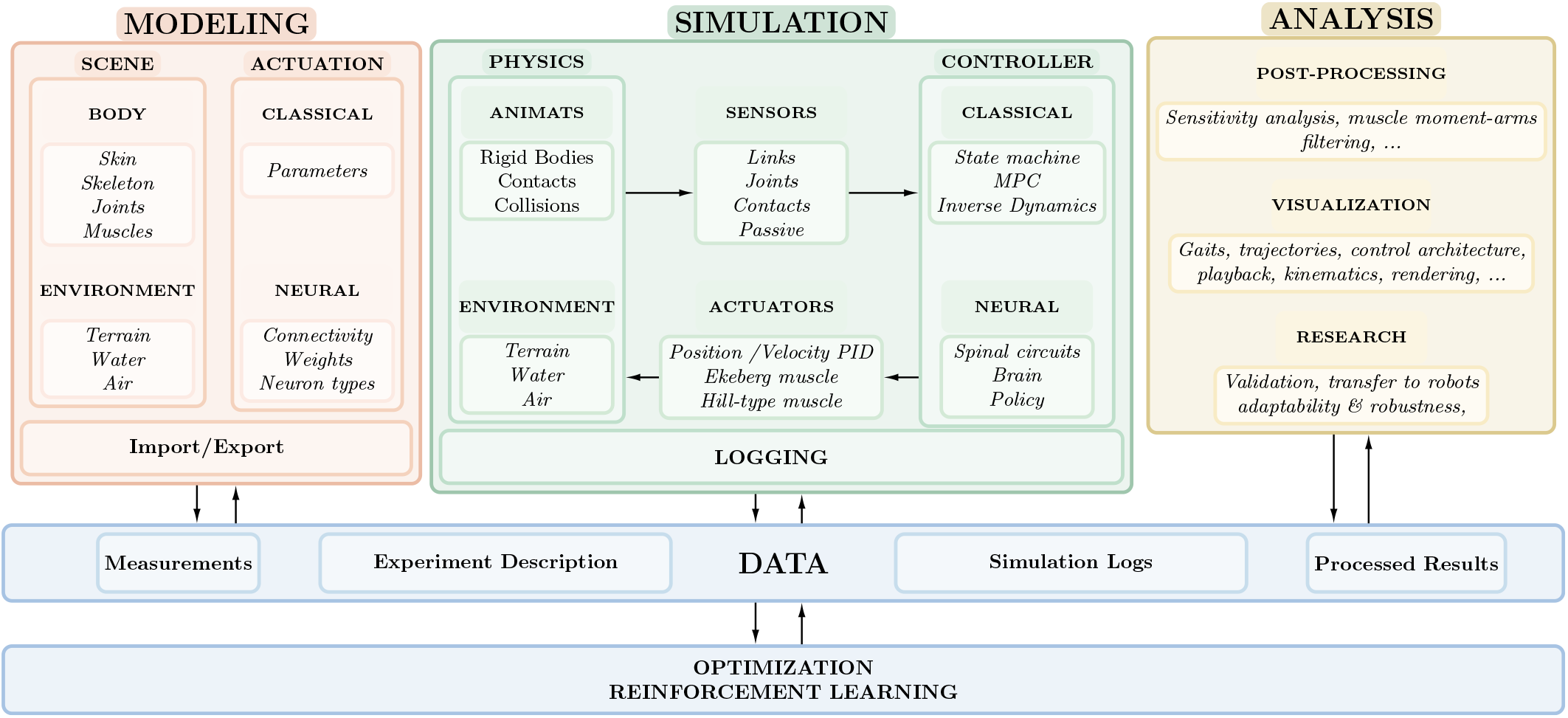
Overview of the modular FARMS architecture. Primary stages of a neuromechanical simulation: modeling, simulation, and analysis. FARMS establishes a standardized data structure that facilitates seamless interoperability across these stages. This structure supports the integration of external libraries, allowing them to interface directly with FARMS and access core neuromechanical simulation data.

- **Data:** At every step of the framework, the information flow is recorded and tracked. This is performed with the data block. Each module can read and write to it. This enables FARMS’ modularity as data can be shared between blocks and can be populated by any independent tool.
- **Modeling:** Two necessities for any neuromechanical simulation are the description of an animal or robot body and its controller for generating motions. The modeling block covers the tools and aspects of generating these sub-blocks.
- **Simulation:** Once both the body and controller are described, the next step consists in running a physics simulation with the controller in the loop. Simulations in FARMS involve integrating (i) a physics engine to simulate articulated rigid bodies and their interactions with the environment (e.g. land or water) and (ii) the control laws to compute the actions to generate movements.
- **Analysis:** Finally, during and after a simulation, the progress of an experiment can be monitored through the collected data. The analysis component deals with performing post-processing, visualizing and replaying simulation data.

Optimization and learning algorithms constitute an additional layer for manipulating each of these components to produce the desired animal or robot behavior.

### 3.1 Data

In FARMS, the reproducibility of simulation workflows is a fundamental requirement. To support this, the complete state of an experiment, including model definitions, simulation parameters, and analysis outputs, can be saved to disk at each stage of the pipeline (modeling, simulation, and analysis). This decoupled approach makes FARMS a modular framework, allowing individual components to operate independently while remaining interoperable. The main package that glues the components of FARMS together is farms_core, which provides consistent data handling across the modeling, simulation, and analysis stages. This unifying approach enhances the modularity of the framework, making it easier to share and reproduce experiments within the research community.

The data block within FARMS consists of three primary types: configuration files, run-time simulation data, and processed simulation data. These data types collectively contain all the necessary information to fully reproduce an experiment (Figure 2).

- **Configuration Files:** These files provide a comprehensive description of the simulation setup required to run experiments accurately. Typically saved in human-readable formats such as Extensible Markup Language (XML) and Yet Another Markup Language (YAML), configuration files detail the animal or robot’s morphology, controller, and initial conditions, as well as environmental parameters such as friction and drag coefficients. They also specify simulation settings like timestep, simulation units, and logging preferences. This ensures that experiments can be replicated precisely, facilitating reproducibility.
- **Run-time Simulation Data:** During the simulation, the data structure provides access to the complete simulation state agnostic to the underlying physics engine, and logging information as specified by the user in the configuration files. This includes detailed records of links (positions, velocities, forces), joints (control commands, forces, limits), contacts (reaction forces), external forces, and muscles (excitations, lengths, forces).
- **Processed Simulation Data:** After the simulation runs, processed data is generated to facilitate analysis and interpretation. The Hierarchical Data Format (HDF5) format is used to store and organize this data, effectively reducing disk space usage [The HDF Group 2023]. The run-time data and simulation results saved to disk share the same structure, enabling researchers to verify exactly how the simulation behaved during post-processing using the same interface as for writing the controller. From this data, derived metrics and visualizations that summarize the experiment’s outcomes can be generated, making it easier to compare results across different simulations and validate hypotheses.

The modularity of FARMS is further enhanced by the seamless interaction between these data blocks. Each module within FARMS can read from and write to the data block using the interface provided by farms_core, allowing different stages of the simulation to be revisited or modified without needing to redo the entire process. For example, a user can adjust parameters in the configuration file and rerun only the simulation stage, using the existing data from the modeling stage.

By systematically saving and organizing data, FARMS not only supports reproducibility but also encourages collaboration and knowledge sharing within the research community. The standardized data formats and comprehensive logging ensure that simulations can be easily shared and reproduced by others, contributing to a more collaborative and transparent research environment.

### 3.2 Modeling

A neuromechanical simulation comprises three core elements: (i) animal or robot bodies, representing both geometric and dynamical properties, (ii) a controller, and (iii) the environment with which the animal or robot interacts. Accurate modeling of each of these elements is a fundamental requirement for neuromechanical simulations. Such models cannot be assumed to be static or readily available, as they must be iteratively refined in response to new measurements or experimental data. Consequently, FARMS is designed to support rapid model reconfiguration, enabling efficient iteration when simulations fail or diverge from experimental observations.

#### 3.2.1 Animat

An experiment can consist of one or more bodies, specifically animals, robots, or both. As models of robots and animals are treated the same way in FARMS, they will be referred to as *animats* hereafter. An animat is defined by several key elements that collectively describe the structural and dynamical properties of its physical body:

- **Rig:** A collection of links and joints that form the structural frame to support and define the entire body.
- **Link:** An individual element in the rig, characterized by its location (position and orientation). It may include an inertial element and some geometries.
- **Joint:** A connection between two links that defines how they interact kinematically and dynamically. Joints can be of different types, including fixed, ball-and-socket, slide, hinge, and a “free joint” for floating bodies. Joints can also be constrained to mimic physical limitations, such as ligaments in animals or mechanical stops in robots, to restrict the range of motion.
- **Inertial element:** An object that describes an individual link’s inertial properties, including mass, centre-of-mass, and inertia.
- **Geometry:** A shape or structure that can have visual and colliding properties interacting with the environment. Each link may possess a set of geometrical elements describing rigid or soft bodies. The visual and colliding properties may differ. Colliding properties sometimes use simplified or primitive shapes that approximate the visual ones to facilitate the collision handling performed by the physics engine.
- **Actuator:** An element that applies forces on the body to generate motion. These can be motors (applying torques or forces at joints) or muscles (applying forces on different links).

These definitions provide the foundation for modeling animats in FARMS, allowing researchers to describe and simulate complex systems accurately.

Even though we represent an animal and a robot under the same description as an animat, the necessary information to define them is obtained differently. Robots are typically designed by humans in Computer Aided Design (CAD) software and usually have a well-defined fixed geometry. However, animal bodies are reconstructed either based on specimen scans [Computerized Tomography (CT) or Magnetic Resonance Imaging (MRI)] or anatomical reference data, and require careful consideration of the intra-animal variability that exists. The scans are then exported as meshes which can be reworked with specialized software, such as Blender [Community 2022]. The meshes are then referenced in the desired simulation description file. Making use of common standards for robot simulation, FARMS relies primarily on the Simulation Description Format (SDF), originally developed by the Open Source Robotics Foundation for the Gazebo robot simulator [Koenig and Howard 2004]. This format is widely used for robotics simulations and contains all the elements required to define the animat models in FARMS. Different views of the information contained within these files are illustrated in Figure 3. While SDF is the main format used for all the models shown in this article, the modularity of FARMS allows integrating support for other formats as well. For example, preliminary support is already present for Unified Robotics Description Format (URDF) and MuJoCo Format (MJCF), two other commonly used formats in robotics simulation originally developed for Gazebo and MujoCo [Todorov et al. 2012] respectively. *OSIM* and *WBT* are some additional examples of frequently used file formats developed for OpenSim [Seth et al. 2011] and Webots [Michel 2004] respectively. While these formats are not officially supported yet, it is possible to convert them to be compatible with FARMS (for example, farms_osim), and further integration remains a future possibility.

**Fig. 3.**
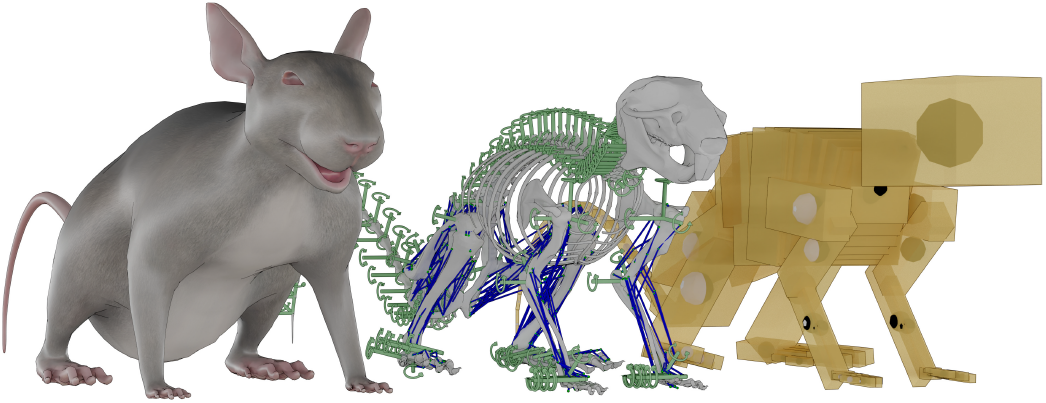
Animat model in FARMS. From left to right visualizes the following elements of a mouse model from Tata Ramalingasetty et al. [2021] (i) skin, (ii) skeleton, joints and muscles, (iii) inertials (inertia tensor, mass and center-of-mass). Segments of skin, skeleton, or simple primitive shapes can be chosen as collision elements to model interactions with the environment.

#### 3.2.2 Blender add-on

Blender [Community 2022] is a full-featured software for 3D modeling, animation, and rendering. It provides a powerful toolkit for creating complex visuals and animations. The farms_blender package is a Blender add-on, providing an interface to users to generate the models and visualize completed simulations, as shown in Figure 4. SDF, URDF and MJCF are based on XML and are typically written in a text editor. The interface provides the ability to set all the links’ poses and geometries, the joints’ poses, as well as the inertials through an interface to facilitate this step. Visual guides can help with validation, as shown in Figure 4a.

**Fig. 4.**
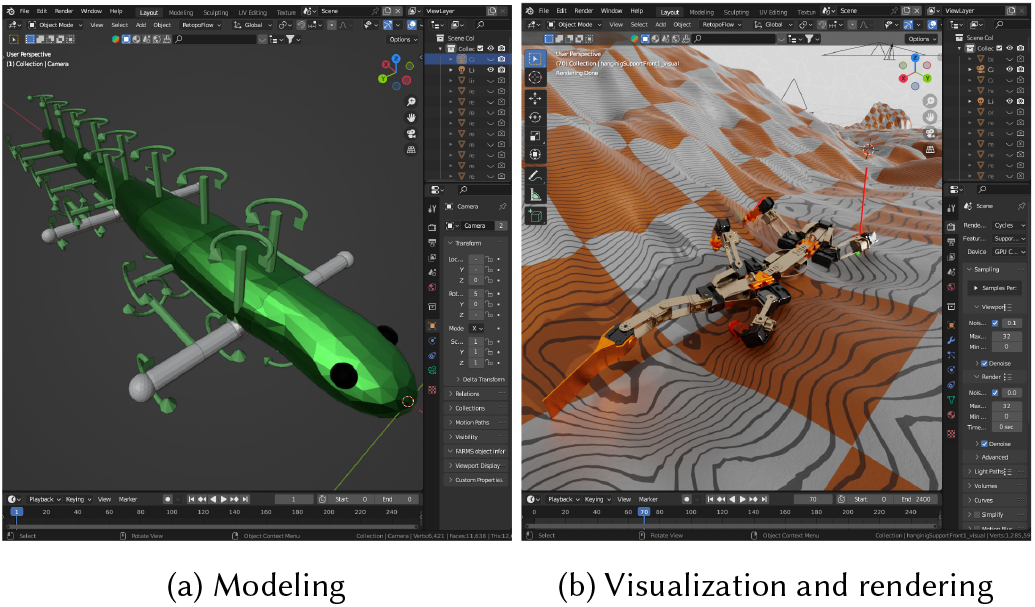
Blender for modeling and rendering using the farms_blender add-on. (a) The modeling example shows the salamander with a display of the joints’ locations and their rotations. (b) The rendering on the right shows a view of a simulation of Krock, a crocodile-like robot, in an uneven terrain after running the simulation and logging data to allow playback within Blender.

The add-on was also developed to fully replay the results of the simulation in Blender, as shown in Figure 4b. This provides a useful interface for reviewing the results of a simulation, showing and overlaying additional information such as interaction forces with the environment, and rendering the simulation results to video for high-quality publications.

#### 3.2.3 Controller

A controller defines the elements that send the commands to the actuators that will ultimately determine how the physical elements move over time. To study animal and robotics movement, several different control strategies are possible. The possible actuation methods currently are (i) model based control and (ii) neural control. Model based control methods are mainly derived from control theory developed for robotics. These include methods like inverse dynamics, state-machines and Model Predictive Control (MPC). Neural circuits with Central Pattern Generators (CPGs)s and sensory feedback loops have proven effective for locomotor control across diverse animals and robots, and are therefore the primary controllers described and presented in the following sections. The use of neural controllers based on deep neural networks using Reinforcement Learning (RL) is also common in recent works. Naturally, the framework grants the ability to implement the other types of controllers as well.

#### 3.2.4 Neural Networks

Neural control typically refers to employing neural network models, whether based on biological principles or machine learning techniques, to generate control signals for movement. This includes biologically inspired CPGs, spiking and non-spiking neural networks, and artificial neural networks. Modeling these systems involves specifying the neuron types, their biophysical or algorithmic properties, and their connectivity, all of which require proper parametrization to generate appropriate behaviors related to complex motor tasks.

Currently, there already exist well-established libraries to build artificial neural networks, such as Pytorch [Paszke et al. 2019] and Tensorflow [Abadi et al. 2016]. Biological networks can be broadly classified into spiking and non-spiking neural networks. For spiking networks, there are open-source libraries such as NEST [Gewaltig and Diesmann 2007], Brian [Stimberg et al. 2019] or Neuron [Hines and Carnevale 1997]. For sparse, non-spiking networks, there is no library to the best of our knowledge that supports modeling and simulation of locomotion networks. As part of FARMS, we developed a platform and simulator agnostic sparse neural network software package called farms_network, which provides commonly used neural models for locomotion circuits. More precisely, it implements standard neural models at different levels of abstraction which have been commonly used in past research, such as phase oscillators [Ijspeert et al. 2007; Kuramoto 1975], Hopf oscillator [Righetti et al. 2006], Matsuoka model [Matsuoka 1985], FitzHugh–Nagumo model [FitzHugh 1961], Leaky-Integrator model [Ijspeert 2001], Hodgkin-Huxley model [Hodgkin and Huxley 1952] and non-spiking activity-based model [Ermentrout 1994]. Figure 5 shows an example of sparse locomotor network based on [Danner et al. 2017], modeled and visualized using farms_network.

**Fig. 5.**
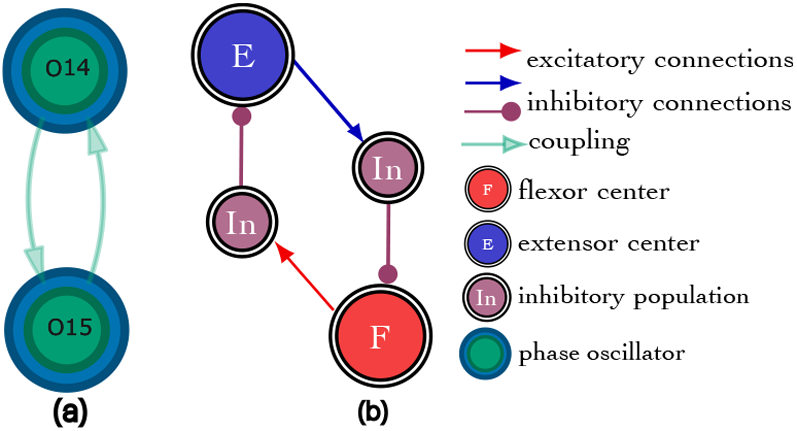
Example of abstract oscillator network models and visualization in FARMS. (a) Coupled phase oscillator network (b) Leaky integrator rhythm generator based on reciprocal inhibition network

While the models implemented in farms_network have no external dependencies, it is straightforward to use the implementations from the well-established libraries for artificial and spiking neural networks, as done in [Pazzaglia et al. 2025].

#### 3.2.5 Actuation

In FARMS, the controller also includes the actuation system, which constitutes the motor dynamics for a robot or the muscle for an animal. In the case of animal locomotion, the biomechanics can play a crucial role in generating the proper gait patterns. Muscles are not a feature often included as part of physics engines as most of them are in practice built for the gaming and robotics communities.

To address this gap, we developed the farms_muscle package, to provide users with different muscle models. Two particular types of muscles conventionally used in literature have been implemented, variations of Hill-type muscle models with options to choose between rigid and compliant tendons in a computationally efficient manner [De Groote et al. 2016; Geyer and Herr 2010; Millard et al. 2013], and the Ekeberg antagonistic muscle pair model [Ekeberg 1993].

As FARMS continues to evolve, the modularity of the framework allows for the future integration of additional muscle models and further refinements to the actuation systems.

#### 3.2.6 Environmen

In FARMS, the environment encompasses both the arena, which defines the terrain geometry, and the interaction properties between the bodies and the surrounding media. The terrain can vary in complexity, ranging from flat surfaces to more rugged landscapes, and can also include obstacles.

Interaction with the terrain is often rigid, as is typical for land-based locomotion. Most physics engines designed for robot simulations primarily support rigid-body interactions, making them well-suited for simulating this type of environment. Depending on the chosen physics engine, FARMS enables flexible interaction models for different terrains. For instance, Bullet [Coumans and Bai 2016] provides robust support for rigid-body dynamics, ensuring accurate simulations of hard surface interactions. Conversely, MuJoCo [Todorov et al. 2012] offers advanced features for soft-body interactions, allowing for more detailed modeling of soft contact dynamics, such as deformations and friction. FARMS provides an interface that allows users to take advantage of these engine-specific capabilities, facilitating accurate neuromechanical simulations across varied terrains and environments.

The framework is also intended to accommodate various complex environmental media, such as water, sand, and air, with each medium characterized by its unique interaction dynamics. This flexibility allows for the simulation of diverse locomotion and interaction scenarios, enabling researchers to study behaviors across a range of physical environments.

### 3.3 Simulation

Once the animat, controller, and environment are fully defined, the next step is to perform numerical simulations. For neuromechanical simulations, this primarily involves iteratively executing the animat controller’s commands and advancing the physics simulation of the entities within the simulated world. During the physics step, the physics constraints must be solved, and the dynamics of the system integrated over time. Leveraging existing physics engines and available frameworks, FARMS provides wrappers that help bridge modalities of the simulation block, illustrated in Figure 2.

#### 3.3.1 Physics engine

The physics engine is the core of a neurome-chanical simulation. The common steps performed by a physics engine are described by Geijtenbeek [2013] as follows:

- **Collision detection** checks if the body intersects with itself or other physical objects in the scene.
- **Forward dynamics simulations** compute the linear and angular accelerations of each simulated body.
- **Numerical integration** updates positions, orientations, and velocities of bodies, based on the previously computed accelerations.

The steps outlined above simulate the articulated rigid bodies and their interactions with rigid terrain. There are several physics engines available, each employing different strategies to perform these three steps. Choosing the appropriate physics engine depends on the required features, as differences in implementation can affect simulation accuracy, stability, and computational cost.

A valuable feature of FARMS is its potential to provide a common interface to multiple physics engines. FARMS currently supports Bullet [Coumans and Bai 2016] with farms_bullet and MuJoCo [Todorov et al. 2012] with farms_mujoco as its primary physics engines. The Bullet and MuJoCo physics engines were mainly selected as primary physics engines in FARMS for their ease of use, extensibility, ability to install on all major desktop platforms, capability to simulate rigid multi-bodies in reduced coordinates for improved stability with many-joint models, and their computational speed when solving simulations of animat locomotion.

Originally, FARMS only supported the Bullet engine, which stood out compared to other engines such as ODE [Smith 2006] for its ease of use through the Python interface and for handling hard collisions reliably without requiring fine-tuning of the contact parameters. While this engine lacked essential features for neuromechanical simulations, such as muscle and fluid models, they could be implemented within FARMS in the farms_bullet package. However, a limitation of Bullet which could not be mitigated is that it implements joint damping explicitly, rather than using an implicit formulation. This led to numerical instabilities for high values of damping, which is an essential feature needed for simulating biomechanical models.

MuJoCo [Todorov et al. 2012] features an implicit implementation of joints damping, facilitating tuning and simulation of muscle-controlled animats. For this reason, the farms_mujoco package was implemented to support MuJoCo and seamlessly migrate certain experiments that required this feature.

MuJoCo also provides some additional beneficial features, such as simple muscle and fluid models. On the other hand, contacts are modeled as soft constraints. While this is a drawback for simulating rigid robot models with hard contacts, the advanced soft-contact model can be desirable to account for the deformation of the skin in animal models.

FARMS supports both headless operation, enabling users to run large batches of simulations efficiently, and a graphical user interface (GUI) that is accessible to beginners as results are visually accessible during execution. Currently, the GUI directly reuses what is provided by Bullet or MuJoCo, depending on which engine was selected by the user for the experiment. This dual capability ensures that FARMS is suitable for both advanced researchers and those new to the field, making it a versatile tool for diverse user groups.

As research requirements evolve and new engines emerge, interfaces to additional simulation platforms and physics engines could be developed in a similar way to the MuJoCo and Bullet integrations.

#### 3.3.2 Simulation step

During each simulation step, FARMS follows a structured process consisting of five sequential stages: Sensors, External forces, Controller, Actuators, Logging, and the Physics engine step. This process repeats iteratively until the simulation reaches a predefined termination condition, e.g. the completion of the simulation duration.

- **Sensors:** At the start of each simulation step in FARMS, the sensor stage collects the state of the simulation as provided by the physics engine for the sensors defined during the modeling phase. This includes exteroceptive signals such as contact forces and ground reaction forces, as well as proprioceptive signals like muscle spindle and Golgi-tendon feedback, joint angles, and velocities. These sensory data points are essential for informing the external forces and controller stages, providing real-time information on the system’s state for subsequent computations. The collected data is stored within a dedicated data interface, allowing efficient access without re-querying the physics engine, thereby minimizing redundant computations and improving performance.
- **External forces:** In this stage, the external forces acting on the animat are defined. These can include user-defined perturbations, such as custom-applied forces or torques, as well as environmental forces derived from predefined properties or dynamic models, such as fluid dynamics or wind resistance. The computation of external forces can depend on sensor data collected in the previous stage, such as velocity, contact forces, or positional information, which inform the calculation of dynamic environmental forces. For instance, drag forces encountered during swimming are computed based on the velocity of each link in contact with the simulated fluid. Using the velocity data from the sensors, FARMS calculates the resulting drag forces, which are then applied as external forces to the animat. As with sensor data, the computed external forces are logged and made available for control and analysis.
- **Controller:** The controller governs the animat’s behavior, determining its actions at each simulation step. Depending on the chosen control approach specified during the modeling phase, this could involve either a model-based control strategy or a neural control system. FARMS provides the controller access to all the sensors and external forces data collected at the previous stage, allowing it to compute the outputs as necessary for the actuators in the next stage. Any user is free to implement their own controller as needed for their application, although the farms_network package is available for facilitating the development of neural controllers. A neural controller usually uses its own solver to integrate the neural dynamics or perform other necessary computations to generate the previously stated outputs, and can operate with an internal timestep to handle stiff equations.
- **Actuators:** The outputs of the controller dictate the actions of the actuators. These outputs include (i) muscle activation signals, (ii) desired joint positions and velocities, and (iii) desired joint torques. FARMS applies the appropriate method for each actuator as defined during the modeling phase, ensuring proper actuation for both robotic and biomechanical systems.
- **Physics engine step:** Once the previous stages are complete, FARMS updates the physics engine with the final outputs from the actuators and external forces. The physics engine then handles the integration step solving physical constraints and updating the dynamics over time. After completing the integration, the stages are repeated for the next iteration, continuing until the simulation reaches its predefined termination.
- **Logging:** For each of the previous stages, FARMS provides data interfaces that automatically log the sensor, external forces, controller, and actuator data. This ensures traceability of the simulation state, providing each stage access to past data, which can be useful for adaptive control and feedback mechanisms. The logged data can also be stored to disk, facilitating post-processing and detailed analysis. This logging feature allows users to revisit and study any point in the simulation, facilitating the extraction of meaningful insights for further research and development.

#### 3.3.3 Extending physics

As part of FARMS, we provide access to both phenomenological and advanced fluid dynamics models. For simplified fluid dynamics, we offer a drag model implemented in both the farms_bullet and farms_mujoco packages, which approximates interactions with fluids through computationally efficient drag forces. For more sophisticated simulations, preliminary support for fluid dynamics based on Smoothed-Particle Hydrodynamics (SPH) methods is available in the farms_sph package. This package integrates the PySPH library [Puri et al. 2013; Ramachandran et al. 2021], enabling two-way coupling between fluid dynamics and rigid-body simulations, similar to approaches described by Angelidis et al. [Angelidis et al. 2022].

Buoyancy forces are also supported and calculated based on the provided volume of each link, generating vertical forces as appropriate. In both the drag model and the SPH-based integration, fluid-induced forces are calculated independently from the rigid-body physics engine. These forces are applied as external forces during each simulation step, with the rigid-body physics engine responsible for integrating the equations of motion. This method ensures compatibility across engines by treating fluid interactions as additional external inputs. While support for sand and air simulations is not yet available in FARMS, a similar methodology could be used to implement these media, leveraging PySPH’s existing capabilities for simulating sand and gas dynamics [Ramachandran et al. 2021]. Development of the SPH integration is still in progress, with plans to extend support to additional SPH models, such as those provided by SPlisHSPlasH [Bender 2017], in future updates.

#### 3.3.4 Optimization

Neuromechanical simulations allow systematic investigation of how parameters governing morphology, control, and environmental physics shape behavior, yet identifying parameter sets that reliably produce desired behaviors remains difficult. Indeed, it can be difficult to measure certain neural or muscular parameters from the animals or impractical to directly test on robot hardware, while simulations can leverage evolutionary algorithms to tune these parameters. For this reason and inspired by Auerbach et al. [2014], we implemented support for evolutionary algorithms in farms_evolution.

Evolutionary algorithms have been employed in a large number of works including studies of lampreys [Ijspeert and Kodjabachian 1999], salamanders [Arreguit et al. 2021; Ijspeert 2001], ants [Guo et al. 2018] and humans [Dzeladini et al. 2014]. More precisely, [Ijspeert 2001; Ijspeert and Kodjabachian 1999] use Genetic Algorithms, [Guo et al. 2018] uses Covariance Matrix Adaptation Evolution Strategy and [Dzeladini et al. 2014] uses Particle Swarm Optimization, all of which optimize a single cost function. The farms_evolution packages leverages the Pymoo library [Blank and Deb 2020], which provides a range of algorithms for single and multi-objective optimization, with or without constraints, for optimizing the parameters of a simulation task.

### 3.4 Analysis

The analysis block plays a crucial role in extracting valuable insights from the simulation data. As shown in Figure 2, the analysis block is seamlessly integrated into the overall architecture. The logging mechanism incorporated in farms_core ensures that all relevant information used to handle the state of the animat at any point of time within the simulation is recorded. With this comprehensive dataset, researchers can for example generate plots of sensor values, evaluate model performance, identify distinct gaits, cluster behaviors, and even replay the results of the simulation to gain deeper understanding without recomputing it.

High-quality visualization tools go a long way in helping researchers in understanding the high dimensional information gathered during simulation. The ability to replay the simulation within a 3D interface is provided via Blender [Community 2022] through farms_blender. This allows simulation replay with overlaid data, providing deeper insights into quantities, such as collision or external forces acting on the animat. By leveraging this capability, scientists can quickly explore various what-if scenarios, optimize their models accordingly and generate high-quality renders for publication purposes.

## 4 Demonstrations

To highlight the features and capabilities of FARMS, we present a number of experiments with different animats, controllers and environments in the sections that follow.

### 4.1 Robots

We first demonstrate a set of locomotion experiments with robots based on eel and salamander-like morphologies with a controller based on oscillator networks (Figure 6).

**Fig. 6.**
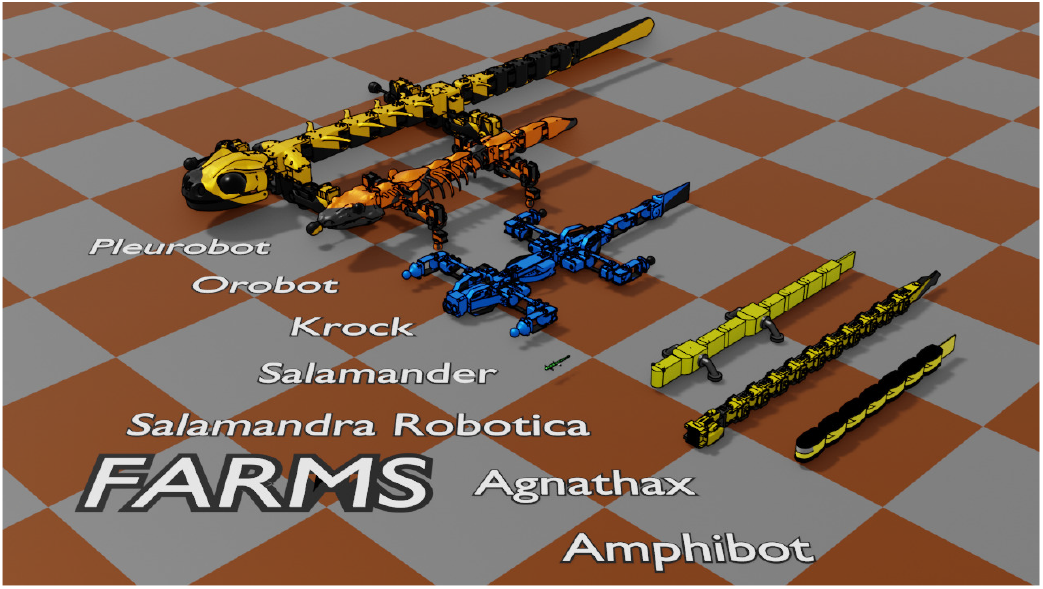
Examples of robots in FARMS. Pleurobot in Karakasiliotis et al. [2016], Orobot in Nyakatura et al. [2019], K-rock in Melo et al. [2017, 2023], Salamandra Robotica II in Crespi et al. [2013]; Ijspeert et al. [2007], Agnathax in Thandiackal et al. [2021], and Amphibot in Crespi et al. [2005]; Crespi and Ijspeert [2006]. The salamander model shown in the center allows for a size comparison between these robots.

We replicated the behaviors presented in Ijspeert et al. [2007], where a salamander-like robot, Salamandra Robotica II [Crespi et al. 2013], is capable of transitioning from swimming in water to walking on land and vice versa. To generate these different locomotor patterns, the controller uses a network of weakly-connected oscillators, based on central pattern generators, and replicated from the work of Ijspeert et al. [2007] which simulates the spinal cord dynamics.

The simulation results obtained for transitions from swimming to walking are shown in Figure 7. This experiment was performed using the MuJoCo physics engine. An arena featuring a ramp and a body of water was created such that the simulated robot could go from one media to another. To demonstrate how FARMS can include custom physics through MuJoCo’s external forces, the forces from the water are applied using a simple drag model, and a buoyancy force allows the robot to swim at the surface of the water given its mass density. The descending drive signals were modulated based on passing a threshold value of contact with the ground such that walking patterns would be generated when the robot is in contact with the ramp. Based on the original work of Ijspeert et al. [2007], we find in Figure 7 that the simulated robot can swim in water, transition to a walking pattern upon ground contact, and walk out of the water.

**Fig. 7.**
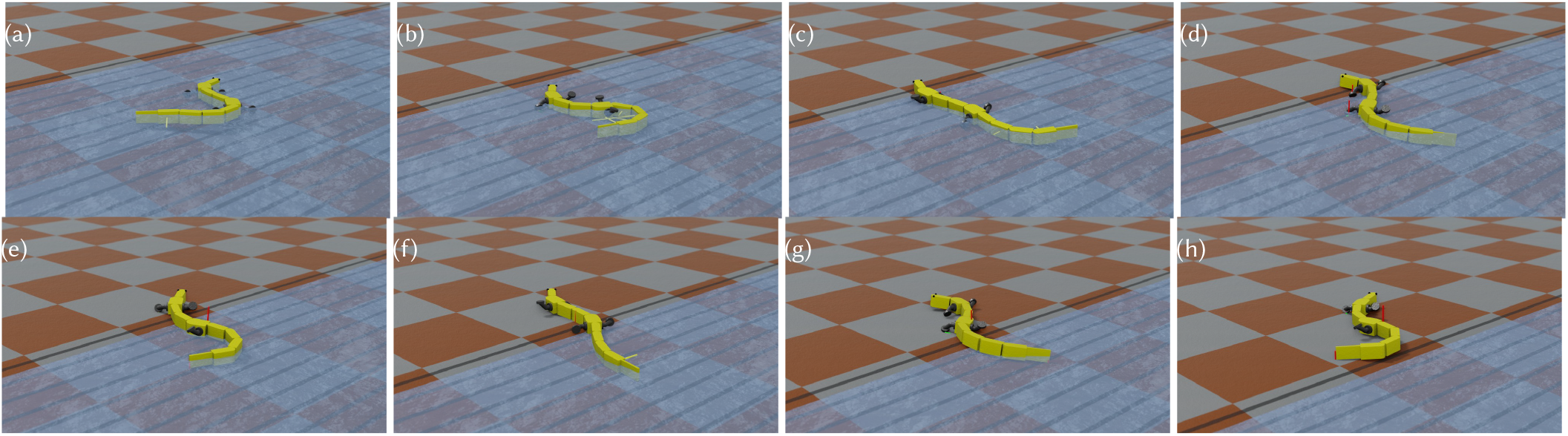
Salamandra Robotica II performing a swim-to-walk transition. Sequence of rendered snapshots shown at different times separated by 1.5 [s]. The robot is swimming from right to left in 7a and comes into contact with the ramp in 7b. In 7c the robot transitions from swimming to walking by changing the descending drive commands and the limbs activate. The trunk oscillators synchronize with the limb oscillators, inducing the body to switch from a travelling wave to a standing wave. From 7c to 7h, the robot has switched to a trotting gait and walks out of the water.

By adapting the oscillator controller to the morphology of the other robots, we show in Figure 8 that walking and swimming gaits can also be obtained with the K-rock [Melo et al. 2017] and Pleurobot [Karakasiliotis et al. 2016] robots models using an adapted network architecture. The particular difference between these robots and Salamandra Robotica II is that they have a higher number of Degrees of Freedoms (DoFs) in the limbs and different numbers of joints within the trunk as well. While still using a double chain of oscillators for the trunk joints, the network is adapted by also using pairs of oscillators for each joint of the limbs.

**Fig. 8.**
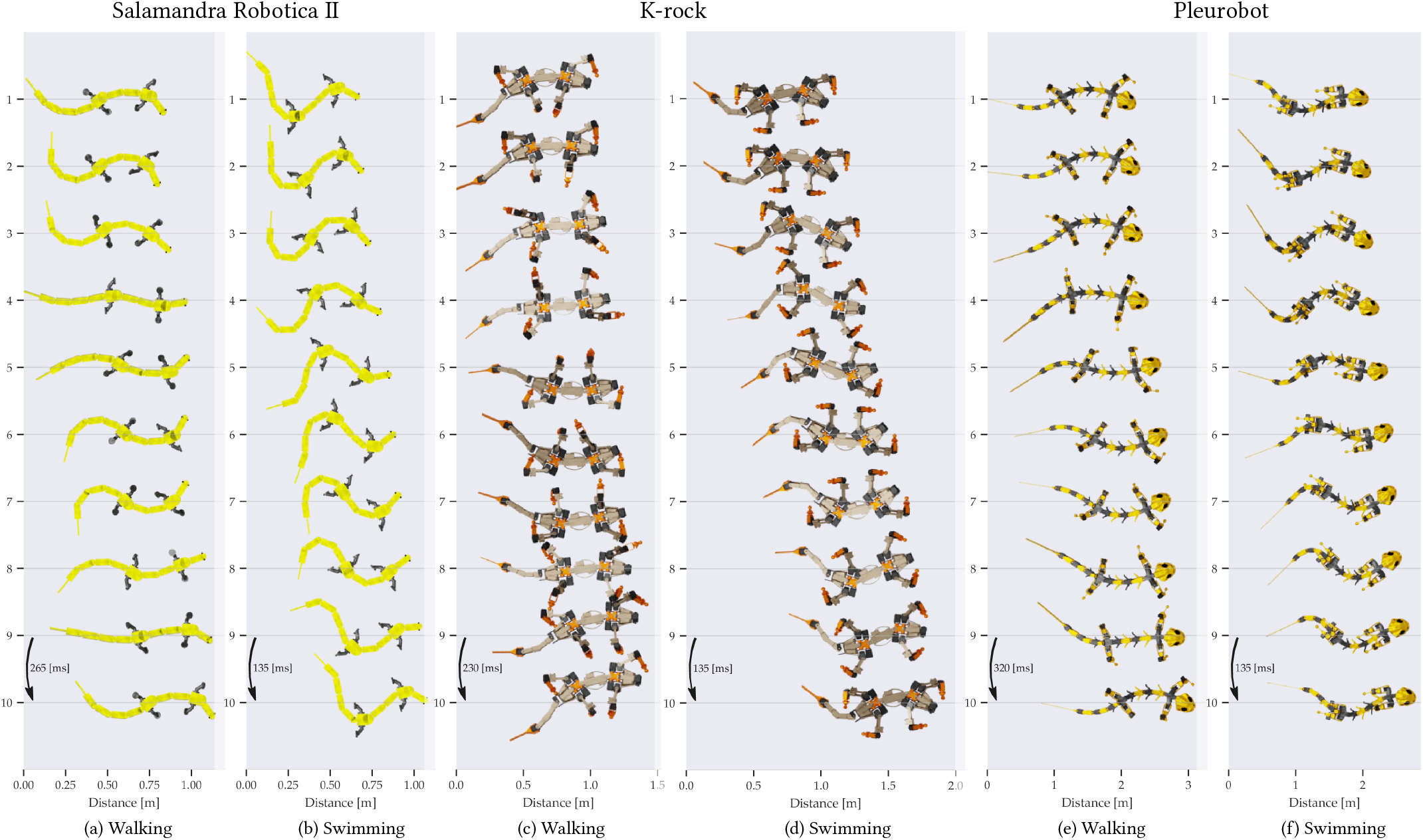
Salamandra Robotica II, K-rock and Pleurobot walking and swimming gaits in simulation. The robots are shown from above with snapshots along time. The spines for each robot exhibit travelling waves which allow them to propel themselves forward in water. A shared controller based on an oscillator network is employed across the robots and systematically adapted to their respective morphologies.

### 4.2 Salamander

Extending the salamander-like robots to animal simulations, a biological salamander model was designed based on the work of Harischandra et al. [2010]. By leveraging the Blender 3D modeling software with the farms_blender package, we provide tools to generate a 3D body shape and segment it into joint-connected rigid bodies, from which masses and inertial properties are automatically computed. Applying these tools yielded an approximate, but comparable representation of the animal. The mass was computed assuming constant density close to water such that the models could float slightly below the surface. The resulting salamander model is 10 cm long and weighs 3 g. It contains 11 body joints allowing the body to bend laterally and 4 DoF for each limb, including 3 DoF at the interface with the body, and one DoF at the elbow/knee.

Similarly to the robots demonstrations, a controller was implemented to showcase walking and swimming using an oscillator network, as shown in Figure 10. However, contrary to the robots presented above in position control, each of the DoF for the salamander is modeled as a revolute joint controlled by a flexion-extension muscle pair model proposed by Ekeberg [1993].

**Fig. 9.**
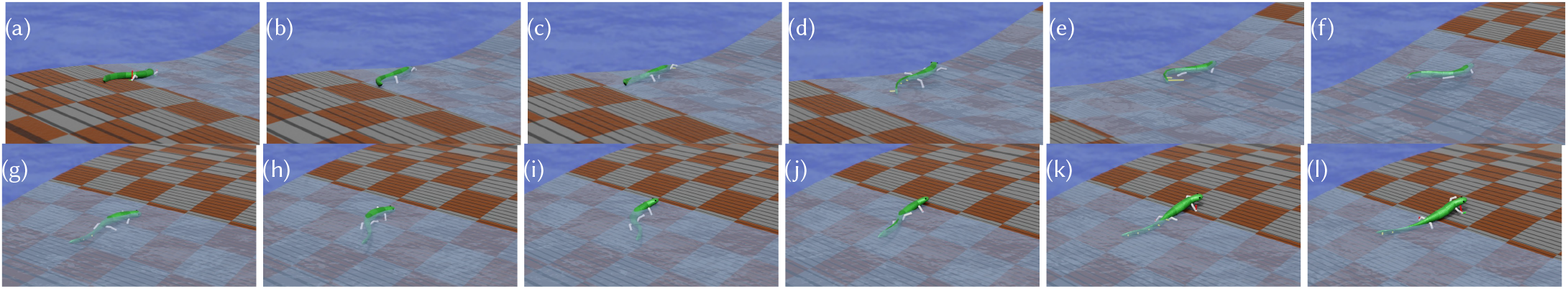
Simulated salamander crossing a river. Sequence of rendered snapshots separated by 1.0 [s]. A salamander is simulated in an arena featuring a river, where a ground created using a heightmap with two banks separated by a river is found. The water has a velocity of 3 [cm/s] flowing from left to right. The salamander is initialized on one side of the river and moves towards the other side. In 9a, the salamander comes into contact with the water, but does not switch to a swimming gait until 9d. From 9d to 9g it swims to the other side pushed by the water flow until it comes into contact with the ground and switches to a walking gait. From 9g to 9l it walks up and out of the water.

**Fig. 10.**
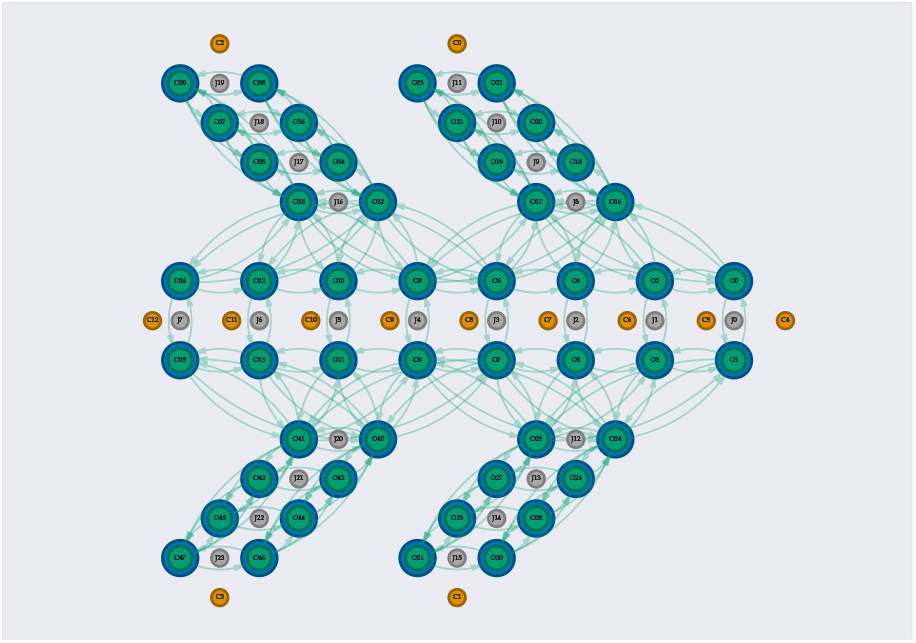
Salamander connectivity for an oscillator network. The network is inspired from the work of Ijspeert et al. [2007]. Here, we explore a connectivity based on weak couplings where oscillators are only connected to neighbors. One of the key differences with Ijspeert et al. [2007] is that the limb oscillators have no direct couplings between them. Although it is known that long-range connections between the limb CPGs exist, here we found that this connectivity was sufficient for the salamander model to achieve walking and swimming patterns, as shown in the demonstrations below.

Just like for the previous demonstrations, this model is also capable of exhibiting both swimming and walking patterns based on a descending drive signal. The main difference is that all the robots in the previous experiments were position controlled, while the joints of the salamander are torque controlled using the muscle implemented in FARMS. In contrast to the Bullet engine, we found that the MuJoCo physics engine was beneficial for properly simulating the dynamics of the Ekeberg muscle model thanks to its implicit implementation of damping.

Figure 9 shows a FARMS simulation of the salamander crossing a river, thereby performing walking-to-swimming and swimming-to-walking transitions. In that experiment, a water current is also present, and the appropriate descending signals are computed to drive the model to follow a path to the other side of the river while switching to the appropriate gait. The passive properties of the muscles are useful for providing compliance to the limbs and permitting the limbs to interact with the ground without predefined trajectories. This model is walking blindly without any knowledge of the surrounding terrain, but shows it can handle it in a robust manner by switching between walking and swimming depending on contact information.

### 4.3 Snake

To extend the salamander results, we also demonstrate how the oscillator network and the passive properties of a muscle model [Ekeberg 1993] can lead to obstacle-based locomotion. A flat arena is set with regularly placed vertical circular pegs with a fixed radius along a grid, as shown in Figure 11. The friction coefficient between the body and the arena is isotropic and set to 0.2 so that the snake model can slip with ease, but cannot advance without pushing on the surrounding pegs. Using a simple double-chained oscillator network for a limbless snake-like body, i.e. the same controller as the salamander but without the limb oscillators, the model is capable of traversing the obstacle configuration. As the model escapes the pegs area, it is unable to move beyond as it cannot push itself against obstacles anymore. Additional experiments for different arena configurations with different values of pegs radius and spacing are shown in the accompanying video. Each of these experiments were performed with no changes in descending signals using the same oscillator network with fixed parameters. This showcases how passive properties of the body can be simulated and interact with the environment with many points of contacts, both with the ground and the pegs, when simulated using the MuJoCo physics engine.

**Fig. 11.**
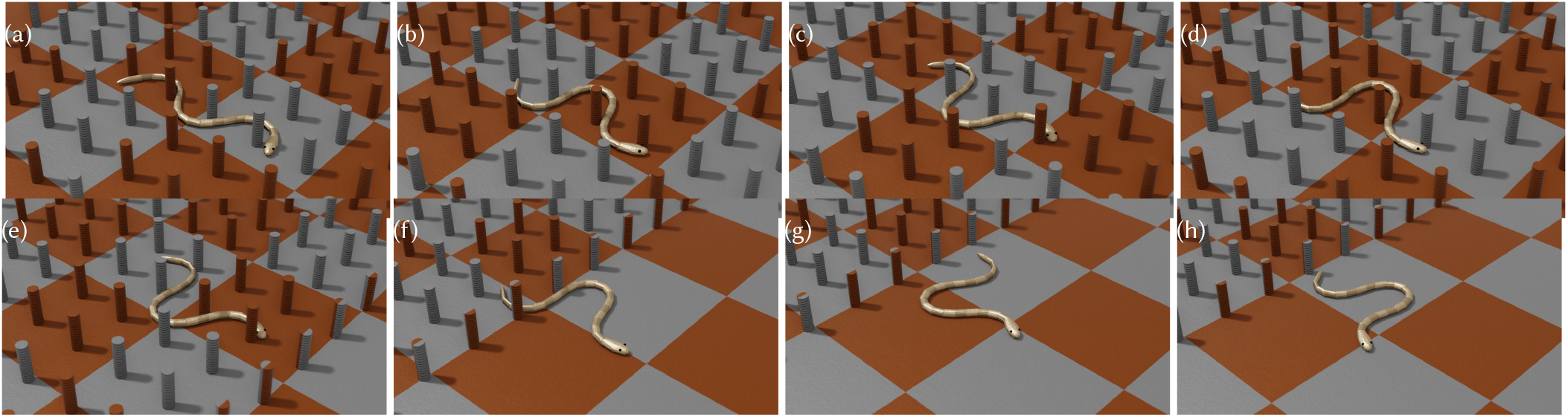
Snake obstacle-aided locomotion in arena 1. Sequence of rendered snapshots separated by 2.5 [s]. The pegs have a diameter of 4 [cm] and are separated by 21 [cm] from centre-to-centre.

### 4.4 Mouse

This section illustrates how FARMS can be used to develop and simulate complex musculoskeletal models from animal data, using a mouse as a case study. A whole-body mouse model for quadrupedal locomotion has been developed and iteratively studied by Tata Ramalingasetty et al. [2021, 2023, 2024].

Constructing Hill-type musculature for articulated skeletons typically requires substantial manual effort, including defining muscle paths across bones and identifying parameters governing muscle force generation. FARMS addresses these challenges by providing an integrated workflow within Blender with the farms_blender plugin that streamlines muscle geometry creation, parameter tuning and estimation. An example of the workflow is shown in Figure 12. Mesh registration (Figure 12 panel 1) tooling is integrated into the plugin to compute affine transformations between corresponding anatomical meshes. This allows muscle attachment points and landmarks defined on one skeletal model to be automatically transferred to another, even when bone geometries differ due to specimen variability. The registration process supports landmark-based alignment for complex meshes and operates directly within the Blender environment, eliminating reliance on external tools. Next, for muscle geometry reconstruction, the plugin provides interactive tools to define multi-point muscle paths directly over skeletal meshes. Intermediate via-points can be added, edited, and visualized in 3D, enabling users to reconstruct anatomically plausible muscle geometry from volumetric scans. This is particularly valuable when no prior musculoskeletal model exists for a given limb or species. Furthermore, the plugin supports automated scaling and estimation of Hill-type muscle parameters. Length-dependent parameters, such as optimal fiber length and tendon slack length, can be adjusted based on skeletal dimensions using a built-in scaling algorithm described in Modenese et al. [2016]. For muscles lacking complete parameter data, we include an extension of the numerical optimization framework described by Manal and Buchanan [2004] to estimate tendon slack length by considering all joint degrees of freedom spanned by the muscle. Together, these features enable construction of detailed musculoskeletal models with minimal manual intervention. In the context of our mouse demonstration, FARMS allowed for efficient transfer of existing muscle data, reconstruction of previously unavailable musculature, and consistent parameterization within a unified environment.

**Fig. 12.**
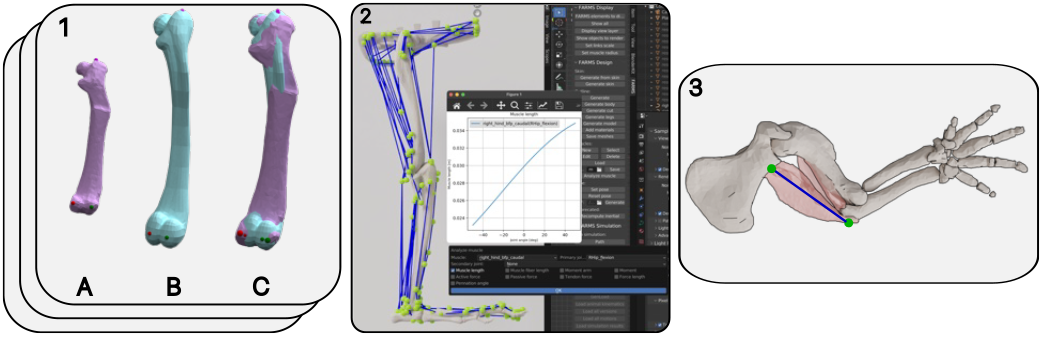
Development of mouse whole-body musucloskeletal model. **Panel 1**: Example of femur bone mesh registration process to transfer muscle attachment landmarks from femur model described by Charles et al. [2016] (1A) to whole-body mouse model described in Tata Ramalingasetty et al. [2021] (1B). 1C validates the successful transfer process. **Panel 2**: Preview of FARMS plugin in Blender to model and analyze musculoskeletal models. The example shows the hindlimb muscusloskeletal model loaded in Blender using farms_blender to highlight muscle modeling and analysis. **Panel 3**: Identification of muscle paths for forelimb muscles based on CT data from DeLaurier et al. [2008].

Finally, Figure 13 shows a simulation of quadrupedal locomotion of a simplified mouse musculoskeletal model controlled by non-spiking activity-based neuronal circuitry described by Tata Ramalingasetty et al. [2023, 2024]. All components of the model—including skeletal morphology, musculature, spinal circuitry, sensory feedback (farms_muscle), and environmental interactions—were modeled and simulated entirely within FARMS. This example demonstrates the framework’s capability to support closed-loop neuromechanical simulations, integrating neural control, musculoskeletal dynamics, and physical interactions in a unified computational workflow.

**Fig. 13.**
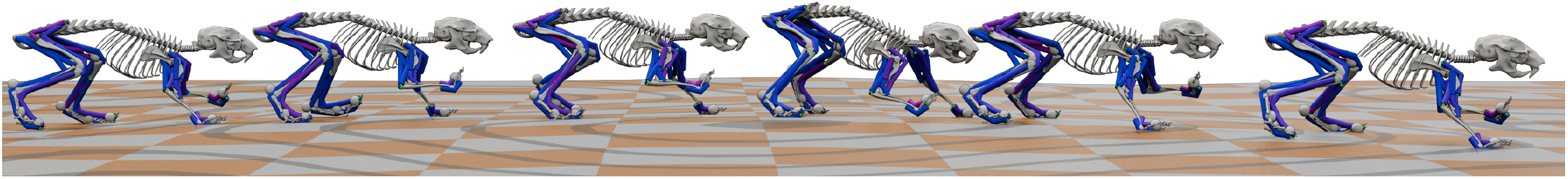
Mouse quadrupedal locomotion in 3D over uneven terrain. A sequence gait snapshots over time of quadrupedal mouse locomotion on uneven terrain controlled by a sensory driven spinal neural model described in Tata Ramalingasetty et al. [2023, 2024].

## 5 Conclusion

Through a series of case studies, we demonstrated that FARMS supports end-to-end, closed-loop neuromechanical simulations, integrating detailed musculoskeletal models, neural control circuits, sensory feedback, environmental interactions, and optimization within a unified framework. These examples span animals and robots, multiple locomotor modes, and diverse environments, illustrating FARMS’ flexibility, scalability, and suitability for studying movement control across biological and engineered systems.

FARMS remains under active development, building on a foundation that already integrates biomechanical, neural, sensory, and learning-based components. Future work will focus on expanding model diversity, environments, and validation, while FARMS continues to evolve as a community-driven, open-source framework supporting reproducible neuromechanical research.

We envision FARMS as a shared, open platform that enables the community to build, compare, and extend neuromechanical models, thereby advancing the study of movement across neuroscience, biomechanics, robotics, and animation.

## 6 Acknowledgments

This research was supported in part by the Human Frontier Science Program (HFSP-RGP0027/2017); in part by the European Research Council (ERC) under the European Union’s Horizon 2020 research (Synergia project, Salamandra); in part by the innovation programme (grant agreement No 951477); European Union’s Horizon 2020 Research and Innovation Program under Agreement 720270917 (SGA1) and Agreement 785907 (SGA2); and in part by the National Institutes of Health (NIH) under Grant R01NS112304, Grant R01NS115900 and in part by 284289 Jekkal Fellowship, Drexel University.

We would like to thank members from Biorobotics Laboratory (BioRob) - EPFL, Neuroengineering Laboratory - EPFL, Danner Lab - Drexel University and Camilla Carta, for valuable feedback on the framework development.

